# Mechanisms involved in the lipid membrane tubulation activity of the pH-responsive MakA protein from *Vibrio cholerae*

**DOI:** 10.64898/2025.12.03.692023

**Authors:** Athar Alam, Abdelbasset Yabrag, Naeem Ullah, Rasmus Sten, Jörgen Åden, Nandita Bodra, Hudson Pace, Marta Bally, Karina Persson, Bernt Eric Uhlin, Sun Nyunt Wai, Aftab Nadeem

## Abstract

Bacterial pore-forming toxins (PFTs) are key virulence factors that disrupt host cell membranes. The *Vibrio cholerae* cytotoxin MakA, a member of the ClyA structural family of α-PFTs, exhibits a unique, pH-dependent ability to induce lipid membrane tubulation. This study elucidates the molecular mechanisms underlying this process. We identify the N-terminal histidine residue (His30) of MakA as a critical pH sensor that is indispensable for membrane tubulation but not required for membrane binding and cell lysis. Substitution of His30 with lysine, MakA^H30K^ , abrogated tubulation without affecting membrane insertion, instead triggering aberrant membrane fusion events. Circular dichroism spectroscopy analysis revealed that the MakA^H30K^ substitution attenuates the robust α-helical transition observed in wild-type MakA upon lipid membrane binding, providing a molecular explanation for its functional defect. Furthermore, we established that membrane cholesterol contributes to MakA-induced tubulation of lipid membranes, as its absence or mutation of a putative cholesterol-binding motif MakA^I236D&I237D^ abolishes both membrane association and tubule formation. A key implication of our work is the mechanistic uncoupling of membrane tubulation from cytolysis, revealing that MakA can perturb membranes via distinct pathways. These findings define a sophisticated activation mechanism where His30 protonation in the acidic host environment and specific cholesterol binding synergistically drive the conformational change necessary for helical oligomerization and membrane tubulation. We discuss our findings from the viewpoint of possible convergent evolution, where bacterial PFTs like MakA partly may mimic ancient membrane remodelers involved in membrane repair pathways, such as ESCRT-III, but for bacterial fitness in host interactions and pathogenesis. Moreover, the present study provides a foundation for possible development of therapeutic strategies aimed at inhibiting these critical interactions.

## Introduction

Bacteria have evolved a remarkable array of strategies to interact with host organisms, and among the most effective virulence factors are proteins that directly compromise cellular membranes. A broad class of these effectors is represented by pore-forming toxins (PFTs), which are secreted as soluble proteins that can undergo conformational rearrangements upon binding to host membranes, eventually inserting into lipid bilayers to generate pores. This process not only disrupts membrane integrity but also triggers a cascade of cellular responses that range from ion imbalance and signaling alterations to cell death [1–4]. PFTs are found across a wide spectrum of bacterial pathogens, both Gram-positive and Gram-negative, and play critical roles in colonization, nutrient acquisition, immune evasion, and microbial competition. Structurally, they are classified into two broad families: α-PFTs, which employ α-helical structures for membrane penetration, and β-PFTs, which form β-barrel pores [3]. Despite structural diversity, the unifying hallmark of PFTs is their capacity to oligomerize on membranes and create aqueous channels that compromise selective permeability.

*Vibrio cholerae*, the etiological agent of cholera, is a paradigmatic example of Gram-negative bacterium that uses an array of toxins to establish infection. The classical virulence determinants are cholera toxin (CT) and the toxin co-regulated pilus (TCP), both of which are essential for the severe diarrheal disease associated with epidemic strains [5]. However, environmental isolates that lack CT and TCP remain pathogenic and have been implicated in secretory diarrhea, wound infections, and septicemia [6, 7]. These strains frequently carry alternative virulence factors, including hemolysins, proteases, RTX toxins, and lipases [7]. Among these, a newly characterized cytotoxin, motility-associated killing factor A (MakA), has emerged as an important fitness factor for *V. cholerae* in its interactions with predators in natural environments and was identified through studies with *Caenorhabditis elegans* and zebrafish infection models [8]. The *makA* gene (vca0883) resides within a five-gene cluster (*makDCBAE*) that is widely conserved among *V. cholerae* isolates [8–10]. Structural studies demonstrated that MakA belongs to the ClyA family of α-PFTs, named after the *Escherichia coli* cytotoxin ClyA [8, 11, 12].

The Mak proteins represent a distinctive subfamily of PFTs. Biochemical analyses revealed that MakA can combine with MakB and MakE to form a tripartite cytolytic complex, while none of these components alone can induce pore formation [9]. This organization resembles other multicomponent α-PFTs, such as YaxAB from *Yersinia enterocolitica* [13], XaxAB from *Xenorhabdus nematophila* [14], and tripartite toxins such as NheABC and HblL1L2B from *Bacillus cereus* [15], AhlABC from *Aeromonas hydrophila* [16], and SmhABC from *Serratia marcescens* [17]. In each case, pore formation depends on the concerted action of all components, often following a precise assembly sequence to maximize cytolytic efficiency [9, 16, 17]. In the case of *V. cholerae*, an equimolar mixture of MakA, MakB, and MakE can assemble into pores in mammalian membranes, underscoring the physiological relevance of this complex [9]. The secretion of MakA, MakB and MakE proteins from *V. cholerae* occurs primarily in a flagellum-dependent manner [8, 9]. Remarkably, *in vitro* studies revealed that MakA alone exerts several independent cellular effects including autophagy modulation, G2/M cell cycle arrest, and apoptosis in cultured epithelial cells [18–20]. Importantly, our recent structural study revealed that MakA possesses a unique, pH-dependent activity distinct from canonical pore formation [21]. Upon exposure to acidic conditions (pH ≤6.5), mimicking the endolysosomal compartment, soluble MakA monomers undergo a large conformational change. The N-terminal head and neck domains refold into transmembrane helices, a process triggered by the protonation of key residues and the unfolding of a protective C-terminal β-tongue. This ’activated’ form inserts into lipid membranes and oligomerizes into unique helical filaments that internalize a thin annular lipid bilayer, rather than forming a classic pore. The oligomerization of MakA drives the formation of high-curvature tubular protrusions that deplete membrane integrity, leading to cytolysis and hemolysis [21]. This pH-dependent mechanism ensures that the MakA activation is precisely targeted within the acidic host environment.

Environmental pH gradients are critical regulators of bacterial virulence, modulating the activity of many membrane-associated proteins. One prominent example is the Phage Shock Protein A (PspA), a bacterial homolog of the Endosomal Sorting Complex Required for Transport-III (ESCRT-III) superfamily. Structural analyses of Synechocystis PspA have shown that its conformational flexibility and ability to assemble into diverse tubular structures underline its pH- and nucleotide-dependent membrane-remodeling activity [22,23]. Notably, PspA possesses intrinsic ATPase activity, and mutations in key residues (R44, E126, and E179) abolish both ATP hydrolysis and membrane remodeling, establishing a direct mechanistic link between nucleotide turnover and function. These findings identify PspA and its homologs as founding members of an ancient ESCRT-III-like superfamily in bacteria, capable of sensing and repairing membrane damage.

The pH-sensing and membrane-remodeling properties of MakA represent a novel extension of this paradigm. Although MakA remains structurally stable across a broad pH range, its membrane tubulation activity is specifically triggered at mildly acidic conditions (pH 5–6.5), mimicking the intestinal environment of the predator model host organism *C. elegans* [21]. Understanding the molecular basis of this pH-dependent activation is therefore crucial to elucidating how *V. cholerae* fine-tunes its virulence mechanisms and potentially exploits ESCRT-III-like strategies to remodel host membranes.

In this study, we investigate the hypothesis that the N-terminal domain of MakA is essential for its membrane perturbation activity under the mildly acidic pH. Building on the observation that this particular MakA activity is pH-dependent, and inspired by the mechanism of nucleotide-sensing in PspA [22, 23], we identified a histidine residue (His30) within the N-terminal domain as a putative pH sensor. We demonstrate that protonation of His30 at acidic pH triggers the conformational rearrangements required for membrane insertion and oligomerization. Substitution of His30 with lysine (MakA^H30K^) abolishes this pH response, effectively eliminating the membrane tubulation activity while preserving membrane binding. These findings reveal that MakA contain a sophisticated pH-sensing mechanism, integrating environmental and lipid cues for its activation. We propose that this mechanism reflects an evolutionarily conserved strategy, reminiscent of ESCRT-III–like membrane remodelers.

## Results

### The histidine residue at the N-terminus of the MakA facilitates tubulation of the target cell membrane

Under mildly acidic conditions, MakA binds to lipid membranes and induces pronounced tubulation [21]. To investigate whether this process requires membrane insertion, we labeled human erythrocytes with the lipophilic dye Dil and incubated them with either unlabeled MakA or Alexa488-labeled MakA (Alexa488-MakA) for 4 hours at pH 6.5, followed by confocal microscopy analysis (**Fig. 1A–B**). The results suggest that MakA inserts into the erythrocyte membrane, as evidenced by the co-localization of Alexa488-MakA with the membrane (**Fig. 1B**). Line profile analysis further validated this co-localization along the tubules (**Fig. 1B**).

**Figure 1:**
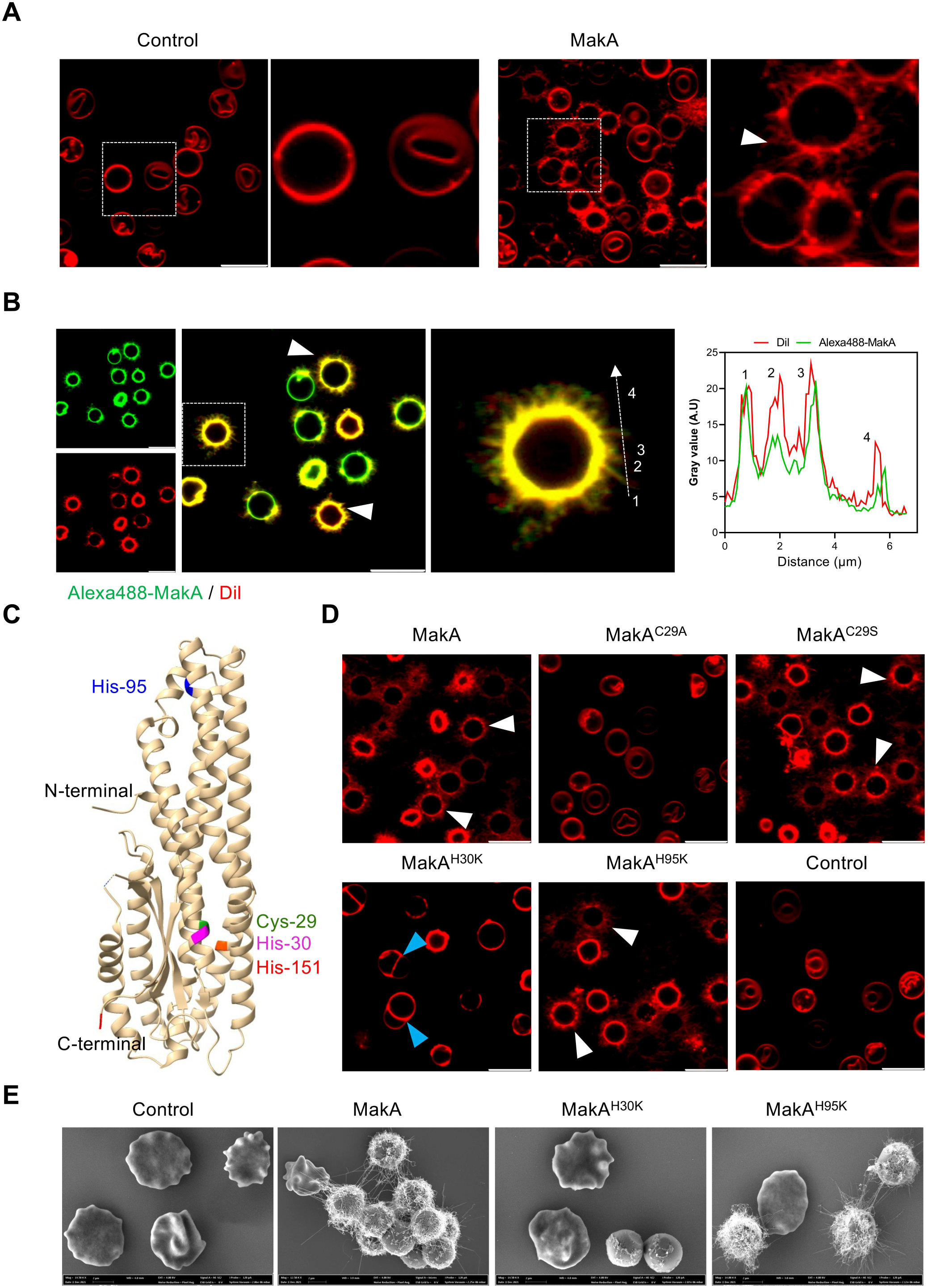
The histidine residue at the N-terminus of MakA facilitates tubulation of the erythrocyte cell membrane. **(A)** Human erythrocytes pre-labeled with the lipophilic dye Dil and suspended in citrate buffer (pH 6.5), were exposed to MakA (1 µM) for 4 hours, followed by live-cell confocal microscopy. White arrowheads indicate tubulation of the cell membrane. **(B)** Cell-associated MakA was visualized using Alexa488-MakA (green). For co-localization experiments, Dil-labeled erythrocytes (red) were incubated with Alexa488-MakA protein (1 µM) for 4 hours at pH 6.5, followed by live-cell confocal microscopy. White arrowheads indicate co-localization of Alexa488-MakA with the Dil-labeled erythrocyte cell membrane. The larger panels show the merged images and an enlargement of one erythrocyte as marked with a square. The line graph (right) shows the fluorescence intensity profile of the corresponding merged image across the dotted white line. The numbers in the image and line plot indicate co-localization of Alexa488MakA with Dil labeled erythrocyte cell membrane across four tubules. **(C)** Crystal structure of MakA monomer (PDB accession no. 6EZV; [8]). Point mutations introduced at the N-terminus of the protein are highlighted in blue (MakA^H95K^), green (MakA^C29A^ or MakA^C29S^), purple (MakA^H30K^), and red (MakA^H195K^). **(D)** Human erythrocytes labeled with Dil (red) were treated with wild-type MakA or the mutant proteins (1 µM) for 4 hours at pH 6.5, followed by live-cell confocal microscopy. White arrowheads indicate tubulation of the erythrocyte cell membrane in response to wild-type MakA or the mutant proteins, while cyan arrowheads in MakA^H30K^ treated cells indicate fusion of erythrocytes. **(E)** Scanning electron microscopy (SEM) images of erythrocytes treated with wild-type or mutant proteins (1 µM, 90 min) in citrate buffer (pH 6.5) are shown. Representative micrographs of erythrocytes indicate the formation of tubular structures on their cell membrane upon exposure to the wild-type MakA or its mutant protein, MakA^H95K^. Membrane perturbation, but lack of tubulation, was observed in the erythrocytes exposed to the mutant protein, MakA^H30K^.

Given the known role of histidine residues in pH sensitive molecular switch [24], we substituted three histidine residues located in the N-terminal and head domains of MakA with lysine, generating the MakA^H30K^, MakA^H95K^, and MakA^H151K^ mutant proteins (**Fig. 1C, and Fig. S1AB**). Based on our recently resolved cryo-EM structural model of membrane-bound MakA, which suggests a homodimeric assembly [21], we also examined the contribution of the adjacent Nterminal cysteine (Cys29) to membrane remodeling. To assess the impact of side-chain polarity and hydrophobicity, Cys29 was substituted with either alanine (MakA^C29A^) or serine (MakA^C29S^) (**Fig. 1C, and Fig. S1A-B**).

All MakA mutants were successfully expressed and purified in soluble form, except MakA^H151K,^ which exhibited poor solubility and was therefore excluded from further analysis. The purity of wild-type and mutant MakA proteins was assessed via SDS-PAGE analysis (**Fig. S1B**). To evaluate membrane remodeling activity, Dil-labeled erythrocytes were incubated with each MakA protein (1 µM, 4 hours at pH 6.5) and analyzed by confocal microscopy (**Fig. 1D**). Both MakA^H95K^ and MakA^C29S^ retained the ability to induce membrane tubulation comparable to wildtype MakA. In contrast, MakA^H30K^ and MakA^C29A^ failed to induce tubulation (**Fig. 1D, and Fig. S2**). Notably, substitution of Cys29 with alanine (MakA^C29A^) completely abolished MakA activity, whereas replacement with serine (MakA^C29S^) preserved function, highlighting the importance of side chain polarity rather than thiol reactivity at this position. Intriguingly, MakA^H30K^ triggered aberrant membrane fusion events, a phenotype not observed in case of wildtype MakA (**Fig. 1D, and Fig. S2**). These findings were corroborated by scanning electron microscopy (SEM), which revealed distinct tubular structures on erythrocyte surfaces treated with wild-type MakA and MakA^H95K^, but not with MakA^H30K^ (**Fig. 1E**). Overall, the data underscores the importance of specific N-terminal residues, particularly His30 and Cys29 were critical for the structural transitions that enable MakA-mediated membrane binding and tubulation.

### Histidine-30 is critical for the pH-sensitive conformational change required for MakA-mediated tubulation

To assess whether substitution of the N-terminal histidine residue affects membrane binding, we incubated Dil-labeled erythrocytes with Alexa488-labeled wild-type MakA, MakA^H30K^, or MakA^H95K^ for 4 hours at pH 6.5 and analyzed the samples by confocal microscopy. Both MakA^H30K^ and MakA^H95K^ mutant proteins were localized to the erythrocyte membrane similarly to wild-type MakA, indicating that these mutations do not impair membrane association (**Fig. 2A**). Despite similar membrane binding, the MakA^H30K^ variant failed to induce tubulation, underscoring that while His30 is dispensable for membrane association, it is vital for pH-dependent conformational activation that drives membrane remodeling. In contrast, the MakA^H95K^ retained tubulation activity comparable to wild-type MakA (**Fig. 2A**).

**Figure 2:**
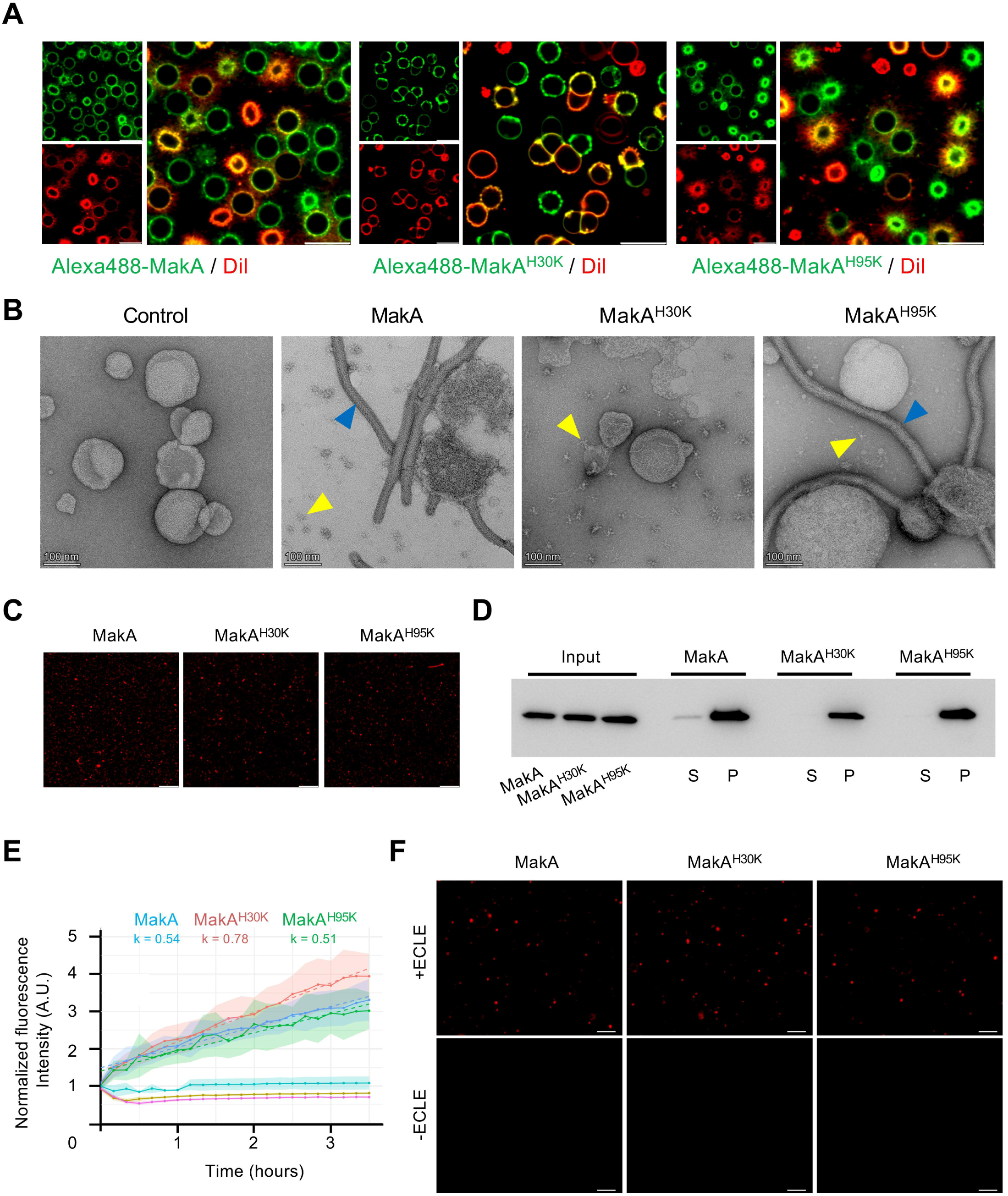
The N-terminus histidine mutant of MakA binds to the lipid membrane but lacks tubulation. **(A)** Cell-associated Alexa488-labeled wild-type MakA or the mutant proteins MakA^H30K^ and MakA^H95K^ were visualized using confocal microscopy. For co-localization experiments, Dillabeled erythrocytes (red) were incubated with the wild-type Alexa488-MakA, and the mutant proteins, Alexa488-MakA^H30K^ or Alexa488-MakA^H95K^ (1 µM at pH 6.5) for 4 hours, followed by live-cell confocal microscopy. The larger panels show merged images. Scale bars = 10 µm. **(B)** Liposomes prepared from epithelial cell lipid extract (ECLE) were treated with the wild-type MakA or the mutant proteins (1 µM at pH 6.5) for 90 min and negatively stained with a 1.5% uranyl acetate solution. Images were captured with transmission electron microscopy (TEM). Blue arrowheads indicate tubular structures, while yellow arrowheads indicate the formation of MakA oligomeric structures present adjacent to or associated with liposomes. Scale bars, 100 nm. **(C)** ECLE liposomes were exposed to wild type Alexa488-MakA or the mutant proteins, Alexa488-MakA^H30K^ or Alexa488-MakA^H95K^ (1 µM at pH 6.5) for 90 min, followed by confocal microscopy. Scale bars = 5 µm. **(D)** Western blot analysis of liposome-bound wild-type MakA and the mutant proteins, MakA^H30K^ or MakA^H95K^ were detected with the anti-MakA antisera. MakA proteins (128 nM) were incubated in citrate buffer (120 mM, pH 6.5) with liposomes prepared from epithelial cell lipid extracts. The pellet indicates liposome bound, while supernatant indicates unbound wild-type MakA or the mutant proteins, MakA^H30K^ and MakA^H95K^. S = supernatant; P = pellet. Equal concentrations of MakA and its mutant proteins were used as an input. **(E)** SparkCyto image analysis for the kinetics of the wild-type MakA, and its histidine mutants (MakA^H30K^ and MakA^H95K^, respectively) binding to liposomes prepared from ECLE. Images were acquired each 10 min for a duration of 3.5 hours. k = linear rate constant per hour. **(F)** Representative SparkCyto images of ECLE liposomes exposed to Alexa488 labeled wild-type MakA, or the mutant (MakA^H30K^ and MakA^H95K^) proteins (1 µM) for 3.5 hours. Scale bars = 20 µm.

To determine whether the loss of tubulation in MakA^H30K^ was specific to the complex erythrocyte membrane environment or reflected a general defect in lipid remodeling, we extended our analysis to reconstituted lipid bilayers. Liposomes prepared from epithelial cell lipid extracts (ECLE) were incubated with wild-type MakA, MakA^H30K^, or MakA^H95K^ (3 μM, 90 min at pH 6.5) and analyzed by transmission electron microscopy (TEM). Consistent with the erythrocyte data, both native MakA and the MakA^H95K^ induced robust liposome tubulation whereas MakA^H30K^ did not (**Fig. 2B**). To ascertain whether this attenuated conformational response in the MakA^H30K^ mutant was due to a loss of lipid binding, we exposed liposomes to native Alexa488-MakA, Alexa488-MakA^H30K^ or Alexa488-MakA^H95K^ (1 µM, 90 min at pH 6.5) and examined them by confocal microscopy. All proteins bound efficiently to liposomes (**Fig. 2C**), indicating that the loss of tubulation by MakA^H30K^ was not due to a defect in membrane binding. Liposome cosedimentation (pull-down) assays further confirmed comparable lipid association across variants (**Fig. 2D**).

We next investigated the binding kinetics of MakA and histidine substitution mutants, i.e., MakA^H30K^ and MakA^H95K^, to liposomes prepared from ECLE using SparkCyto imaging system (**Fig. 2E-F**). Data analysis indicates that MakA and those mutants (MakA^H30K^ and MakA^H95K^) have similar binding profiles for liposomes prepared from ECLE. Importantly, there was no significant difference in linear rate constant per hour among the mutant proteins and wild-type MakA (**Fig. 2E**). Collectively, these results demonstrate that His30 was not required for membrane binding but appeared indispensable for the pH-sensitive conformational change that enables MakA-driven membrane tubulation.

### Histidine-30 is dispensable for membrane lysis but critical for membrane fusion

A hallmark of many membrane-active proteins, such as PFTs and antimicrobial peptides, is their ability to undergo a conformational transition from a disordered state in solution to an α-helical structure upon membrane association [25, 26]. To investigate whether the MakA^H30K^ or MakA^H95K^ substitution mutations would have an impact on conformational change compared to the wild-type MakA, we analyzed their secondary structures at pH 6.5 using circular dichroism (CD) spectroscopy in the presence and absence of liposomes. The far-UV CD spectra of all three proteins in solution were nearly identical, indicating that the substitution mutations do not cause major structural alterations under these conditions (**Fig. 3A**). Upon the addition of liposomes, all proteins exhibited the characteristic increase in α-helical content, evidenced by a significant intensification of the minima at 208 nm and 222 nm (**Fig. 3B**). Quantitative comparison of the spectral changes (**Table S1**) revealed that the MakA^H95K^ mutant protein underwent conformational reorganization comparable to the wild-type MakA. In contrast, the MakA^H30K^ mutant displayed a markedly attenuated helical transition, suggesting it fails to adopt the fully active, membrane-bound conformation required for efficient tubule formation. This structural defect provides a plausible molecular explanation for its inability to induce membrane tubulation. We have previously demonstrated that MakA induces hemolysis of erythrocytes when tested at acidic pH [21]. However, a potential mechanistic link between MakA-induced membrane tubulation and cell lysis has remained unclear. To investigate whether membrane tubulation is a prerequisite for MakA-mediated lysis, we examined the hemolytic activity of wild-type MakA and the histidine substitution mutant proteins (MakA^H30K^ and MakA^H95K^) using both confocal microscopy and quantitative hemolysis assays (**Fig. 3C-D**). Confocal microscopy of Dil-labeled erythrocytes exposed to MakA proteins for 4 hours at pH 6.5 revealed comparable levels of membrane disruption across all conditions (**Fig. 3C**). Notably, despite its inability to induce membrane tubulation, the MakA^H30K^ variant still caused erythrocyte lysis, indicating that tubulation is an essential prerequisite for membrane rupture. This observation was supported by quantitative hemolysis assays, which showed similar levels of hemoglobin release from the erythrocytes treated with native MakA and its histidine mutants, MakA^H95K^ and MakA^H30K^ respectively (**Fig. 3D).** Interestingly, while MakA^H30K^ failed to induce tubulation (**Fig. 1D and Fig. 2A-B**), it promoted membrane fusion events between erythrocytes, a phenotype not observed in cells treated with native MakA and MakA^H95K^. These fusion events, however, did not correlate with increased hemolysis, suggesting that fusion and lysis represent mechanistically distinct outcomes of MakA-membrane interactions. Together, these results demonstrate that while His30 is not required for MakA-mediated membrane lysis, it plays a critical role in promoting the conformational and structural rearrangements necessary for membrane fusion and tubulation.

**Figure 3:**
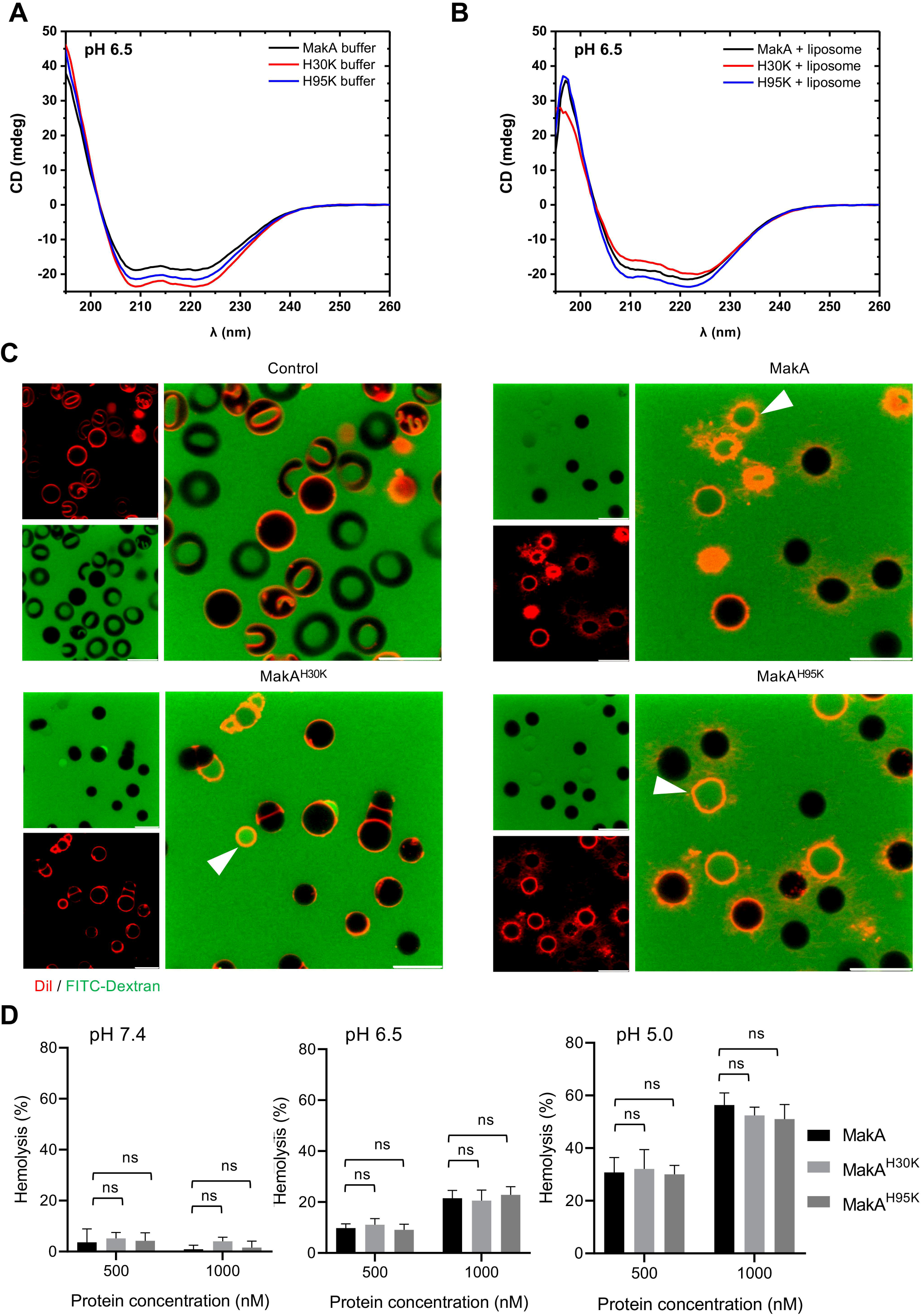
MakA-induced lysis of human erythrocytes does not depend on the pH-sensing His30 residue. Far-UV circular dichroism (CD) spectra of wild-type MakA and the mutant (MakA^H30K^ or MakA^H95K^) proteins were recorded in the absence **(A)** or presence **(B)** of liposomes prepared from ECLE. CD spectra were obtained in citrate buffer (5 mM, pH 6.5) using 3 µM protein. The absorption intensity measured from buffer-only controls was subtracted to correct background absorption. **(C)** Human erythrocytes pre-labeled with Dil were suspended in citrate buffer (pH 6.5) and exposed to wild-type MakA or the mutant (MakA^H30K^ or MakA^H95K^) proteins (1 µM at pH 6.5) for 4 hours, followed by live-cell confocal microscopy. FITC-Dextran (green, 1 mg/mL) was added to the extracellular medium to assess cell permeability. White arrowheads indicate erythrocyte lysis. **(D)** Human erythrocytes suspended in citrate buffer (pH 6.5) were treated with increasing concentration of wild-type MakA or its mutant (MakA^H30K^ or MakA^H95K^) proteins for 5 hours. Hemolytic activity was quantified by measuring released hemoglobin and normalized to erythrocytes treated with 0.1% Triton X-100 (set as 100% lysis). Data are representative of two independent experiments; bar graphs show means ± s.d. Significance from the replicates was determined using two-way ANOVA with Tukey’s multiple comparison test. ns = not significant.

### Cholesterol is promoting MakA-induced membrane tubulation

Cellular membranes are composed of diverse lipid species, among which cholesterol is a key structural component in modulating membrane properties of erythrocytes and other mammalian cells [27]. Our previous work demonstrated that MakA binds to synthetic lipid membranes (SLMs) of defined composition, including cholesterol, and induces pronounced membrane tubulation [21]. Notably, we also observed tubulation induced by MakA at acidic pH on liposomes prepared with lipids extracted from *C. elegans* or *E. coli*, organisms that lack cholesterol [21]. To further delineate the specific lipid requirements underlying this process, we systematically tested MakA activity on SLM variants differing in lipid composition. TEM analysis of SLM liposomes exposed to MakA (3 µM, 90 min at pH 6.5) revealed robust tubulation occurred in all tested SLM variants except those lacking cholesterol (**Fig. 4A**). This finding indicates that cholesterol can be a critical component required for MakA-induced membrane remodeling in mammalian cells. To determine whether cholesterol is required for MakA membrane association, we incubated SLM liposomes containing or lacking cholesterol with Alexa568-MakA (3 μM, 4 hours at pH 6.5) and analyzed binding by confocal microscopy. The results showed a marked reduction in MakA binding and liposome aggregation in the absence of cholesterol (**Fig. 4B**), suggesting that cholesterol facilitates MakA attachment and subsequent structural reorganization of the membrane.

**Figure 4:**
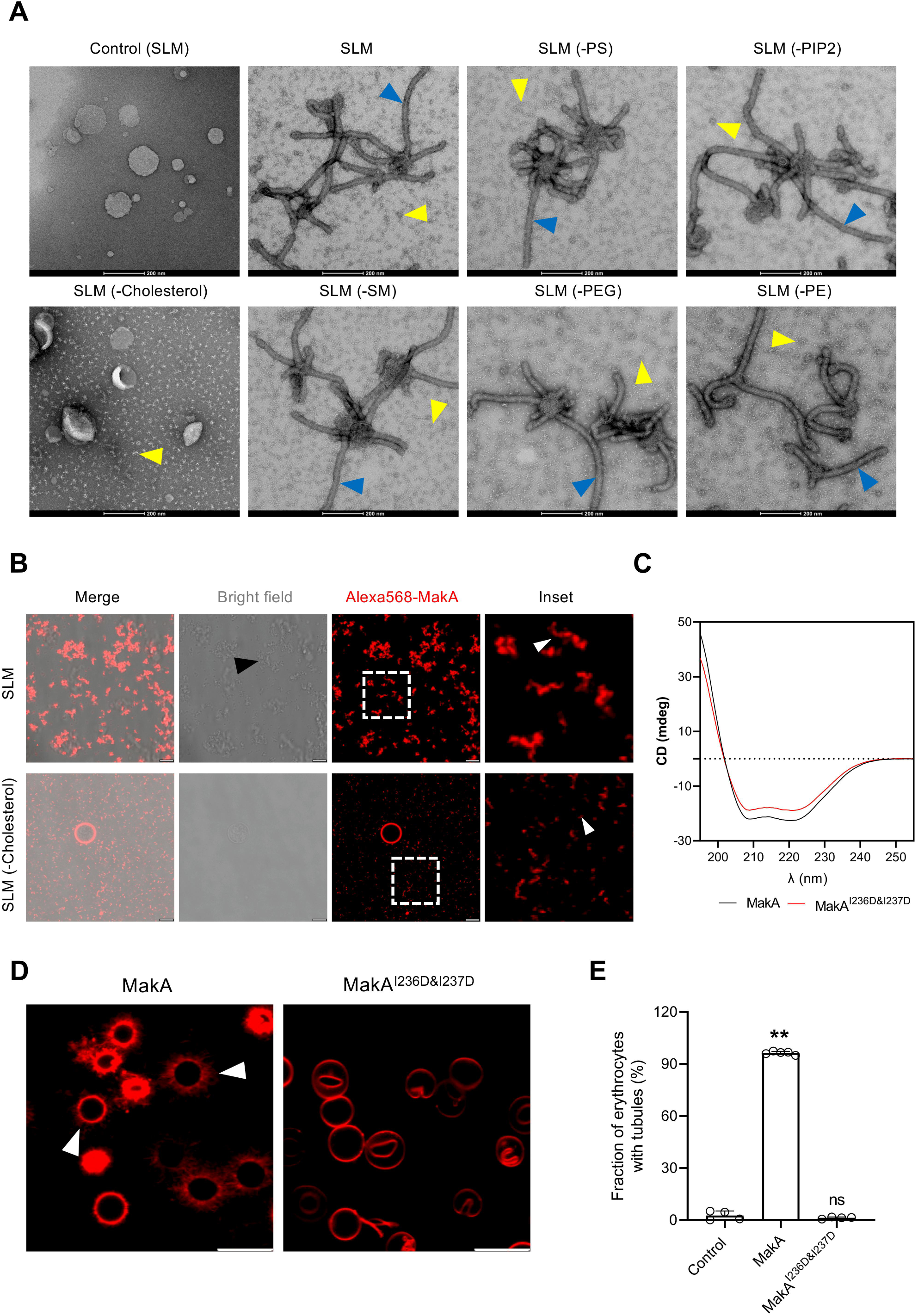
Cholesterol contributes to induction of lipid membrane tubulation in response to MakA **(A)** TEM images of synthetic lipid mixture (SLM) liposomes after incubation with MakA (3 µM, 90 min) in citrate buffer (pH 6.5). Robust tubulation was observed in all SLM variants except those lacking cholesterol. Blue arrowheads indicate tubular structures, while yellow arrowheads indicate the formation of MakA oligomeric structures present nearby or on liposomes. Scale bars, 200 nm. **(B)** Confocal microscopy of Alexa568-labeled MakA (3 µM, 4 hours at pH 6.5) incubated with SLM liposomes containing or lacking cholesterol. Reduced fluorescence and liposome aggregation (shown with black arrowhead) were observed in the absence of cholesterol, indicating impaired MakA binding. White arrowheads indicate liposome-bound MakA. Scale bars, 5 µm. **(C)** Far-UV circular dichroism (CD) spectra of wild-type MakA and the mutant (MakA^I236D&I237D^) protein were recorded in phosphate buffer (pH 6.5) using 3 µM protein. The absorption intensity measured from buffer-only controls was subtracted to correct for background. **(D)** Confocal microscopy analyses of human erythrocytes labeled with the lipophilic dye Dil (red) exposed to wild-type MakA or the mutant (MakA^I236D&I237D^) protein (1 µM) in citrate buffer (pH 6.5) for 4 hours. White arrowheads indicate membrane tubulation of erythrocytes. **(E)** The histogram indicates quantification of number of erythrocytes with tubules (n = 156 to 214 cells / field of view) as shown in panel D. The data point represents quantification of number of erythrocytes from 4 to 5 fields of view (180 x 180 μm^2^), collected from two independent experiments. Data is presented as mean ± s.e.m.; one-way analysis of variance (ANOVA) with Sidak’s multiple comparisons test. **p ≤ 0.01, ns = not significant.

To further dissect the molecular basis of this cholesterol dependence, we examined the MakA^I236D&I237D^ mutant protein in which the hydrophobic residues Ile236 and Ile237 were substituted with aspartic acid (**Fig. S1A**). Those residues were previously implicated in cholesterol recognition by MakA [28]. Circular dichroism (CD) spectroscopy revealed minimal changes in secondary structure among MakA and the MakA^I236D&I237D^ mutant (**Fig. 4C**) indicating that the overall protein fold remained intact. Functionally, however, the MakA^I236D&I237D^ mutant protein failed to induce tubulation in both erythrocyte membranes and SLM liposomes, in contrast to wild-type MakA (**Fig. 4D-E**). Collectively, these results demonstrate that cholesterol strongly promoted MakA-mediated membrane tubulation on the tested SLM liposomes. The requirement for cholesterol likely reflects its dual role in promoting MakA binding and facilitating the conformational rearrangements necessary for membrane deformation.

## Discussion

The MakA protein is the first α-PFT protein family member from *V. cholerae* shown to possess the ability to induce lipid membrane tubulation [21]. In this study, we aimed to identify key molecular determinants underlying the membrane-active properties of *V. cholerae* MakA. The Nterminus (2-42 residues) of MakA has been previously implicated in membrane interaction [18]; however, the specific amino acid residues mediating this interaction remained undefined. Our present study revealed that the N-terminal histidine residue (His30) is indispensable for acid sensing and membrane tubulation and that cholesterol serves as a cofactor required for MakA-induced membrane remodeling of synthetic lipid membranes. Furthermore, we found that His30 was not required for hemolysis activity, suggesting that membrane tubulation and lysis at pH 6.5 arise from distinct mechanistic pathways. Substitution of His30 with lysine (MakA^H30K^) abrogated tubulation without affecting membrane binding, as evidenced by multiple complementary approaches, including confocal microscopy, transmission- and scanning-electron microscopy, and liposome co-sedimentation assays. Histidine residues are well known as molecular pH sensors due to the proximity of their side-chain pKa to physiological pH, allowing them to undergo reversible protonation and deprotonation in response to environmental acidification [24]. The importance of histidine-mediated conformational rearrangements in membrane-active toxins has been demonstrated for several bacterial PFTs, where protonation drives structural transitions required for membrane insertion and oligomerization [29–32]. Our results extend this paradigm to MakA by demonstrating that His30 is specifically required for the transition from a membrane-bound to a tubulation-competent conformation. The MakA^H30K^ mutant protein, while capable of binding and partially inserting into membranes, fails to undergo the robust α-helical transition observed in wild-type MakA upon lipid interaction, as shown by CD spectroscopy. This attenuated conformational response likely underlies its inability to induce membrane tubulation. Interestingly, the MakA^H95K^ mutant protein, in which a histidine residue close to the head domain is substituted, retained both membrane binding and tubulation activity, indicating that the N-terminal His-30 plays a unique and non-redundant role in acid sensing and structural activation. This specificity is further supported by our observation that the MakA^H30K^ mutant protein, but not MakA^H95K^, induces membrane fusion events, suggesting that His30 may also influence the mode of membrane remodeling. The ability of MakA to induce different membrane perturbances depending on its structural state and the lipid environment underscores the versatility of this protein to affect cell membranes via diverse molecular mechanisms. Mammalian cellular membranes are composed of diverse lipid species, with cholesterol serving as a key structural determinant of membrane rigidity, curvature, and microdomain organization [27]. Our data demonstrate that MakA activity is strongly promoted by cholesterol for effective membrane binding and tubulation. Absence of cholesterol abrogated MakA association with synthetic lipid membranes and markedly reduced liposome aggregation, indicating that cholesterol directly facilitates MakA–membrane interactions. Furthermore, substitution of two hydrophobic residues (Ile236 and Ile237), previously implicated in cholesterol recognition, with negatively charged aspartic acids MakA^I236D&I237D^ variant [28] abolished MakA’s ability to bind membranes and induce tubulation. This finding supports the notion that MakA recognizes and exploits cholesterol-rich domains, such as lipid rafts, as platforms for membrane remodeling. Similar cholesterol-promoted membrane sculpting has been reported for eukaryotic proteins such as caveolin-1 (Cav-1), which binds to cholesterol-rich domains to generate positive curvature and induce tubulation [4]. Thus, MakA may mimic or exploit host-like mechanisms of membrane remodeling to perturb cellular integrity during predator/host interactions.

Our findings demonstrate that MakA can induce tubulation in lipid membranes of liposomes derived from diverse biological sources, including human epithelial cell extracts, *E. coli* total lipid extracts, and *C. elegans* total lipid extracts. This broad activity suggests that MakA recognizes conserved physicochemical membrane features rather than some specific lipid composition(s). In liposomes prepared from a synthetic lipid mixture (SLM), however, membrane tubulation was strictly dependent on the presence of cholesterol, highlighting a critical role for sterols in facilitating MakA-mediated membrane remodeling. This sterol dependence is consistent with evidence that cholesterol-driven changes in membrane order, rigidity, and organization facilitate pore-forming toxin (PFT) binding, insertion, and oligomeric assembly [33]. The *E. coli* lipid extracts are primarily composed of phosphatidyl ethanolamine (PE), phosphatidylglycerol (PG), and cardiolipin (CL), with PE as the most abundant neutral lipid and anionic PG and CL conferring a net negative charge [34, 35]. The *E. coli* total lipid extract may support tubulation through mechanisms independent of sterols. Notably, CL accumulates in highly curved membrane regions, and its conical shape favors curvature, presumably lowering the energetic barrier for MakA-induced tubule formation and potentially enabling direct interactions that promote membrane remodeling [36, 37]. In contrast, *C. elegans* membranes, dominated by PC and PE glycerophospholipids, may contain very low levels of cholesterol [38–40]. The observation that MakA can still induce tubulation in these membranes suggests that even trace amounts of sterols may suffice for MakA activity. Such low-level sterols may derive from dietary uptake under laboratory conditions, where *C. elegans* relies entirely on exogenous sources for sterol acquisition [41]. This highlights the intriguing possibility that MakA can exploit minimal sterol content to initiate membrane curvature, pointing to a high sterol sensitivity or a cooperative mechanism with other lipid species. MakA-induced tubulation in these membranes suggests that MakA may interact with a broader spectrum of sterols than previously recognized, although differences in sterol structure could influence efficiency or mechanistic details. These observations emphasize the need to dissect sterol-specific contributions to MakA activity and to understand whether MakA distinguishes between sterols in facilitating membrane insertion, oligomerization, or curvature induction. Together, these findings highlight MakA’s versatility in remodeling membranes with diverse lipid compositions and point to key unanswered questions about sterol recognition and its implications for PFT function, membrane biophysics, and predator/host–pathogen interactions across evolutionary distances.

MakA-induced membrane remodeling resembles processes mediated by integral and peripheral membrane proteins that sculpt membranes through hydrophobic insertion or scaffolding mechanisms. For instance, integral membrane proteins such as reticulons and A17, encoded by poxvirus, generate high-membrane-curvature structures like vesicles and tubules through hydrophobic domains with intrinsic shapes, such as conical or inverted conical transmembrane domains [42, 43]. Reticulons, with their short hairpin transmembrane domains, occupy more space in the outer leaflet, adopting an inverted cone shape that drives curvature, while A17 forms crescent-shaped viral structures with tubules as small as 25 nm in diameter [42, 43]. Similarly, FAM134B, an endoplasmic reticulum (ER)-resident protein, uses two wedge-shaped transmembrane hairpin domains to generate high membrane curvature in the ER, facilitating vesicle formation for phagosomal engulfment [44, 45]. Similar to MakA, peripherin-2/rds and the influenza virus M2 protein induces tubulovesicular membrane foci by inserting its amphipathic αhelices (AH) in cholesterol rich lipid membranes [46]. MakA appears to follow a comparable strategy: upon acid-induced activation and cholesterol binding, its amphipathic α-helices reorganize into oligomeric filaments that drive local curvature and membrane protrusion. Moreover, MakA’s membrane remodeling activity draws intriguing parallels with the eukaryotic ESCRT-III complex, which plays a critical role in membrane repair and scission [47]. While MakA, as a bacterial protein has presumably evolved to disrupt membranes in a pH-dependent manner to promote *V. cholerae* fitness against predatory organisms, ESCRT-III functions to restore membrane integrity to maintain cellular homeostasis, such as in multivesicular body (MVB) formation, cytokinesis, and wound repair [22]. Despite their seemingly opposing roles, both systems rely on dynamic α-helical assemblies capable of inducing extreme curvature. MakA induces high-curvature tubulation (∼25-50 nm diameter tubules) through pH-dependent oligomerization into helical filaments, promoted by cholesterol binding and His30-mediated conformational changes. ESCRT-III subunits (e.g., CHMP proteins) polymerize into spirals to constrict membranes during intraluminal vesicle formation or wound repair. Although MakA lacks sequence homology to ESCRT-III components, the structural resemblance of their helical assemblies suggests convergent evolution, whereby MakA may exploit similar biophysical principles for destructive remodeling. Notably, host cells have been shown to recruit ESCRT-III machinery to repair pores generated by bacterial PFTs [23], raising the possibility that MakA-induced damage may trigger compensatory ESCRT-dependent repair responses.

A striking observation from the current study is that the His30 residue, while essential for tubulation, was not required for MakA-mediated hemolytic activity at acidic pH. The MakA^H30K^ variant, despite its inability to induce tubulation, retains hemolytic activity, as demonstrated by confocal microscopy and quantitative hemolysis assays. This observation demonstrated that membrane tubulation and lysis are mechanistically distinct processes. Tubulation likely reflects a specialized structural state of MakA optimized for membrane remodeling, whereas lysis results from alternative, perhaps less ordered, modes of membrane disruption. This functional bifurcation is reminiscent of cholesterol-dependent cytolysins (CDCs), which can exist in distinct pre-pore and pore-forming states, or antimicrobial peptides that induce a spectrum of membrane perturbation, from transient leakage to catastrophic rupture, depending on concentration and lipid composition [48, 49]. Our data thus suggests that MakA possesses multiple membrane-active configurations that enable it to exert diverse biophysical effects on host membranes.

In summary, this study reveals that the His30 residue acts as a molecular pH sensor mediating MakA’s conformational activation, while cholesterol serves as a structural cofactor that promotes membrane binding and tubulation. Together, these factors orchestrate MakA’s ability to remodel, fuse, and rupture membranes through distinct but interrelated mechanisms. These findings provide fundamental insights into how this protein from *V. cholerae* can be employed for manipulation of mammalian cell membranes and establish a conceptual framework for exploring bacterial protein–membrane interactions for possible biotechnological applications.

## Material and Methods

### Purification of native MakA and its variants

The cloning, expression, and purification strategy for native MakA has been described previously [8]. To engineer the substitution mutants at the N-terminal of MakA (designated MakA^C29A^, MakA^C29S^ MakA^H30K^, MakA^H95K^, and MakA^H151K^) or the head domain of MakA^I236D&I237D^, the wild-type MakA plasmid served as the template for site-directed mutagenesis using the QuikChange XL-II kit (Stratagene). The desired substitution mutation was confirmed by DNA sequencing. The primers used for this mutagenesis were: MakA_C29A-F 5’-tgt gtt taa aat ggc atg ggc ctg cgc tgt gat gac tgt g-3’, MakA_C29A-R 5’- cac agt cat cac agc gca ggc cca tgc cat ttt aaa cac a-3’; MakA_C29S-F 5’-cag tca tca cag cgc aga gcc atg cca ttt taa ac-3’, MakA_C29S-R 5’-gtt taa aat ggc atg gct ctg cgc tgt gat gac tg-3’,; MakA_H30K_R 5’-aat tgt gtg ttt aaa atg gcc ttg cac tgc gct gtg atg act g-3’, MakA_H30K_F 5’-cag tca tca cag cgc agt gca agg cca ttt taa aca cac aat t3’; MakA_H95K_R 5’-gtt gga tcc gct tta tat aac tcc tta atg gca tcg att gat gct tga a-3’,

MakA_H95K_F 5’-ttc aag cat caa tcg atg cca tta agg agt tat ata aag cgg atc caa c-3’; MakA_H151K_R 5’-gca cca ttg act aag tca tcc ttt gct gct tgc att ttc acc c-3’, MakA_H151K_F 5’ggg tga aaa tgc aag cag caa agg atg act tag tca atg gtg c-3’. For protein expression, *E. coli* BL21(DE3) LysS cells (Novagen) were transformed with plasmids encoding either the wildtype or mutant MakA proteins. Cultures were grown to an OD of approximately 0.8, at which point protein expression was induced with 1 mM IPTG. Following induction, cells were harvested by centrifugation at 4 °C. Protein purification was carried out using Ni-NTA affinity chromatography. The N-terminal His -tag was cleaved using TEV protease, and the mixture was passed through a second Ni-NTA column to remove uncleaved protein and protease. The resulting protein was concentrated using Amicon Ultra-15 centrifugal filters with a 10 kDa molecular weight cutoff (Millipore) and further purified by size-exclusion chromatography on a HiLoad 16/60 Superdex 200 pg column (GE Healthcare) equilibrated in PBS (pH 7.4). Protein purity was assessed by SDS-PAGE. Purification of MakA^H151K^ was not performed due to its inability to remain in soluble form.

Fluorescent labeling of both wild-type MakA and mutant proteins were performed using the Alexa Fluor 488 or Alexa Fluor 568 Protein Labeling Kit (Thermo Fisher Scientific), following the manufacturer’s instruction.

### Circular dichroism spectroscopy

Far-UV circular dichroism (CD) spectra of wild-type MakA and the mutant proteins (MakA^H30K^, MakA^H95K^, MakA^I236D,I237D^) were recorded, with or without ECLE liposome complexes, using a Jasco J-720 spectropolarimeter (Jasco, Japan) at 25 °C. Protein samples (3 µM) and ECLE liposomes (1 mg/mL) were incubated overnight at 25°C in either citrate (pH 6.5) or phosphate buffer (pH 6.5). Measurements were conducted in a 0.1 cm path length quartz cuvette with a bandwidth of 2 nm. Spectra were collected over the 190–260 nm range, averaged across five scans, and corrected by subtracting the corresponding buffer baseline. All measurements were performed under identical conditions to ensure comparability across variants. The reproducibility of spectra was confirmed by repeated scans, and no significant baseline drift was observed.

### Human erythrocyte hemolysis assay

Freshly prepared human erythrocytes (0.25%) suspended in citrate buffer (120 mM sodium citrate, pH adjusted to 7.4, 6.5 and 5.0) were incubated with increasing concentrations of MakA or MakA^H30K^ or MakA^H95K^ mutant proteins in Eppendorf tubes for 5 hours at 37°C. Following incubation, samples were centrifuged at (500 × *g*) for 10 min, and the supernatants were collected. Hemoglobin release, indicative of red blood cell lysis, was quantified spectrophotometrically by measuring absorbance at 415 nm. Hemolysis induced by MakA or its mutants was normalized to erythrocytes treated with 0.1% Triton X-100 (set as 100% lysis). Data are presented as the percentage of total hemolysis relative to the Triton X-100 treated cells.

### Extraction of epithelial cell lipids for liposome binding assays

Lipids were extracted by the Folch method (Folch et al., 1957) from 10 × 150 cm^2^ confluent flasks of HCT8 cells. Briefly, the lipid extracts dissolved in chloroform were dried to a thin film under a gentle nitrogen stream, yielding approximately 12 mg of total lipid. The dried lipid film (5 or 10 mg/mL) was hydrated in one of the following buffers: HEPES buffer (10 mM HEPES, 150 mM NaCl, pH 7.4), or citrate buffer (120 mM citrate buffer, pH 6.5). The resulting lipid suspensions were extruded through polycarbonate membranes (0.1 μm) using an Avanti MiniExtruder (Avanti Polar Lipids, Alabaster, AL, USA).

The liposome pull-down assay was performed as previously described [21]. Briefly, the liposome suspension was diluted five-fold in freshly prepared binding buffer (120 mM sodium citrate, pH 6.5) and incubated with 128 nM of either native MakA or histidine mutants, MakA^H30K^ or MakA^H95K^. The mixtures were incubated at 37°C for 60 min and then centrifuged at 21,000 ×g for 30 min at room temperature. The pellets were washed two to three times with their respective binding buffers to minimize background binding. Samples were resuspended in buffer, boiled, and analyzed by SDS-PAGE. Proteins were then transferred to nitrocellulose membranes, blocked with skim milk, and probed with anti-MakA primary antibodies (1:10,000 dilution in 5% skimmed milk) overnight at 4°C. After washing, membranes were incubated with HRPconjugated secondary antibodies, and signals were visualized using a chemiluminescent substrate. Imaging was performed using an ImageQuant LAS 4000 system.

### Confocal Microscopy

Freshly prepared human erythrocytes (0.25%) in suspended citrate buffer (120 mM sodium citrate, pH adjusted to 6.5) were prelabelled with the liphophilic dye Dil (5 μM). The labeled erythrocytes were then treated with wild-type or mutant MakA proteins (1 µM) in 18-well chamber slides (µ-Slide, ibidi) for 4 hours at 37°C in a 5% CO_2_ incubator.

For binding assay with Alexa488-MakA, Alexa488-MakA^H30K^, or Alexa488-MakA^H95K^, freshly Dil-labeled (5 μM) human erythrocytes (0.25% in citrate buffer, pH 6.5) were loaded into an eight-well chamber slide (µ-Slide, ibidi) and incubated with Alexa488-MakA (1 µM) for 4 hours at 37°C in a 5% CO_2_ incubator.

For FITC-dextran uptake assays, DiI-labeled erythrocytes (0.25%) were exposed to wild-type MakA or mutant (MakA^H30K^ and MakA^H95K^) proteins (1 µM) in citrate buffer (pH 6.5) for 4 hours in 18-well chamber slides (µ-Slide, ibidi), followed by the addition of FITC-Dextran (1 mg/mL) for 30 min at 37°C.

Liposomes prepared from SLM including or lacking cholesterol (SLM-cholesterol) were incubated with Alexa568-MakA (1 µM) for 4 hours at 37°C prior to imaging. All samples were visualized using a Leica SP8 inverted confocal system (Leica Microsystems) equipped with an HC PL APO 63×/1.40 oil immersion lens.

### Scanning Electron Microscopy

Freshly prepared human erythrocytes (0.25%) were treated with MakA, MakA^H30K^, or MakA^H95K^ proteins (1 µM, 90 min) in citrate buffer (120 mM, pH 6.5). Samples were fixed with a fixative solution (1% glutaraldehyde +0.1 M CaCo buffer +3 mM MgCl_2_), followed by two washes with buffer (0.1 M CaCo buffer +2.5% sucrose +3 mM MgCl_2_). The fixed cells were allowed to sediment onto poly-L-lysine-coated coverslips for 1 hour and subsequently dehydrated in a series of graded ethanol solutions in the microwave. Samples were then subjected to critical point drying (Leica EM300) and coated with a 2 nm platinum layer using a Quorum Q150T ES sputter coater. Imaging was performed using a Carl Zeiss Merlin field-emission scanning electron microscope equipped with an in-chamber (ETD) secondary electron detector at an accelerating voltage of 5 kV and a probe current of 150 pA.

### Transmission Electron Microscopy (TEM)

Negative staining for liposomes prepared from epithelial cell lipid extracts (ECLE) or synthetic lipid mixture (SLM) was performed on glow-discharged copper grids (300 mesh) coated with a thin carbon film (Ted Pella, Redding, CA) as previously described [21]. Briefly, ECLE or SLM of various lipid compositions were incubated with wild-type MakA, or mutant (MakA^H30K^ or MakA^H95K^ ) proteins (3 μM) for 90 min in citrate buffer (pH 6.5) before preparation for TEM. A 3 μL aliquot of each sample were applied to the grids, washed twice with Milli-Q water (MQ) water, and stained with 1.5% uranyl acetate solution (EMS, Hatfield, PA), followed by a final wash with Milli-Q water. Grids were examined using Talos L120C TEM, operated at 120 kV. Images were captured with a Ceta 16M CCD camera using TEM Image & Analysis software ver. 4.17 (FEI, Eindhoven, The Netherlands).

### SparkCyto image analysis

The kinetics of wild-type MakA and mutant proteins, MakA^H30K^ and MakA^H95K^, were investigated using the SparkCyto imaging system. Alexa568-MakA, Alexa568-MakA^H30K^ and Alexa568-MakA^H95K^ (1 μM each), were incubated in the presence or absence of liposomes prepared from ECLE. Images were acquired every 10 minutes for a total duration of 3.5 hours, with and without ECLE liposomes. The fluorescence intensity from the images was quantified as integrated intensity using Fiji ImageJ [50]. A total of four spots, each measuring 583931 μm², were quantified from each image. After pooling conditions between experimental repeats, mean blank subtraction, normalization against time zero, the linear rate constants (k)/hour were calculated.

### Statistical analysis

Results from replicates are presented as mean ± s.d. Statistical significance between groups was determined using one-way or two-way ANOVA, performed in GraphPad Prism. Significance levels were defined as p < 0.05 (*), p < 0.01 (**), and ns = not significant.

## Acknowledgements

A.N. received support from the Swedish Research Council (2022-04779) and the Kempe Foundations (JCSMK23-0138). We acknowledge the facilities and technical assistance of the Umeå Core Facility Electron Microscopy (UCEM) and the Biochemical Imaging Center (BICU), Umeå University, a part of the National Microscopy Infrastructure NMI (VR-RFI 201600968 and VR-RFI 2019–00217). We acknowledge the Protein Expression and Purification facility (PEP) at Umeå University for construct design and cloning.

## Author Contributions

All co-authors contributed to the design of experiments, data analysis, interpretation, manuscript revisions, and agree on the final contents of the manuscript.

## Author Information

Correspondence and requests should be addressed to A.N.

